# Polar Gini Curve: a Technique to Discover Single-cell Biomarker Using 2D Visual Information

**DOI:** 10.1101/2020.03.04.977140

**Authors:** Thanh Minh Nguyen, Jacob John Jeevan, Nuo Xu, Jake Chen

## Abstract

In this work, we design the Polar Gini Curve (PGC) technique, which combines the gene expression and the 2D embedded visual information to detect biomarkers from single-cell data. Theoretically, a Polar Gini Curve characterizes the shape and ‘evenness’ of cell-point distribution of cell-point set. To quantify whether a gene could be a marker in a cell cluster, we can combine two Polar Gini Curves: one drawn upon the cell-points expressing the gene, and the other drawn upon all cell-points in the cluster. We hypothesize that the closers these two curves are, the more likely the gene would be cluster markers. We demonstrate the framework in several simulation case-studies. Applying our framework in analyzing neonatal mouse heart single-cell data, the detected biomarkers may characterize novel subtypes of cardiac muscle cells. The source code and data for PGC could be found at https://figshare.com/projects/Polar_Gini_Curve/76749.

## Introduction

Discovering biomarkers from the single-cell gene expression data is an interesting yet challenging problem [1]. Compared to the well-established bulk gene expression data, the expression distribution in single-cell is significantly more heterogeneous [2–4]. Therefore, as shown in [5, 6], the bulk-analysis strategies [7, 8] achieve low sensitivity in detecting markers. In addition, as embedding [9–11] and clustering [12–14] are the essential components in many single-cell expression analytical pipelines [15, 16], the biomarker detection techniques would need to tackle the challenges and errors from embedding and clustering [17, 18].

From the statistical point of view, there are two different directions among the current state-of-the-art methods in solving the single-cell biomarker discovery problem. The first direction is using non-parametric approaches [19]. Non-parametric approaches do not attempt to construct the model characterizing the gene expression distribution [20]. They do not require too many prior assumptions about the expression data. Therefore, in theory, they could be applied in most of the heterogeneous scenarios in single-cell expression. For example, Seurat [16] and the SINCERA [21] pipelines use the Mann–Whitney test [22]. The disadvantages of non-parametric approaches include lacking the point-estimator (for example, we could not tell how much of fold-change when comparing the expressions of the same gene in two populations) and the lower true positive rate [5, 6]. On the other hand, the parametric approaches model the underlying expression distribution. For example, [23] applies Bayesian statistics, Monocle2 [11, 24] and MAST [2] apply different linear models, and [25] applies the Poisson models to single-cell differential expression analysis. The parametric approaches, compared to the non-parametric ones, are significantly more sensitive [5, 6], especially in detecting markers in small cell-cluster since they may require less number of cell-samples. However, these approaches assume that the gene expression distribution has specific shapes; therefore, these approaches tend to have higher false-positive rates.

In this work, we developed a new framework based on the novel idea of integrating expression and the embedded visual information of single-cell data into one metric to identify biomarkers. This idea has been successfully implemented in spatial single-cell data, in which the visualization space reflects the relative position of the cells in a tissue image [26, 27]. In this framework, we decided to take advantage of cluster shape and cell-point distribution from the 2D visual space. Our strategy was to project the single-cell 2D cluster onto multiple angle-axes to explore all viewing angles of the cluster. On each ‘viewing angle’, we captured the visual distribution using the Gini coefficient [28]. Together, for each set of points in 2D, we constructed a Polar Gini Curve (PGC) from the correspondent between viewing angle and Gini coefficient. We hypothesized that for the marker gene, its expressing cell set should have its PGC close to the PGC computed from the whole cluster cell-set. We demonstrated the framework in several simulation case-studies. Applying our framework in analyzing neonatal mouse heart single-cell data [29], the detected biomarkers may characterize novel subtypes of cardiac muscle cells. We named the framework PGC-RSMD (Polar Gini Curve – Root Mean Square Deviation). The source code and dataset, including supplemental data, used in this manuscript could be found in https://figshare.com/projects/Polar_Gini_Curve/76749.

## Material and Method

### Computing PGC-RSMD for one gene in one cluster

**Figure 1** demonstrates the workflow to compute PGC-RSMD for one gene in a cell cluster from the single-cell expression data. Our approach used the 2D embedding [9] and clustering results from single-cell expression data as the input. Starting from the 2D *x*-*y* embedding space, for an arbitrary angle θ, the pipeline projects the x-y coordinate [30] *for every cell-point* onto the θ-axis (*z* score)

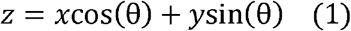

**Figure 1.**
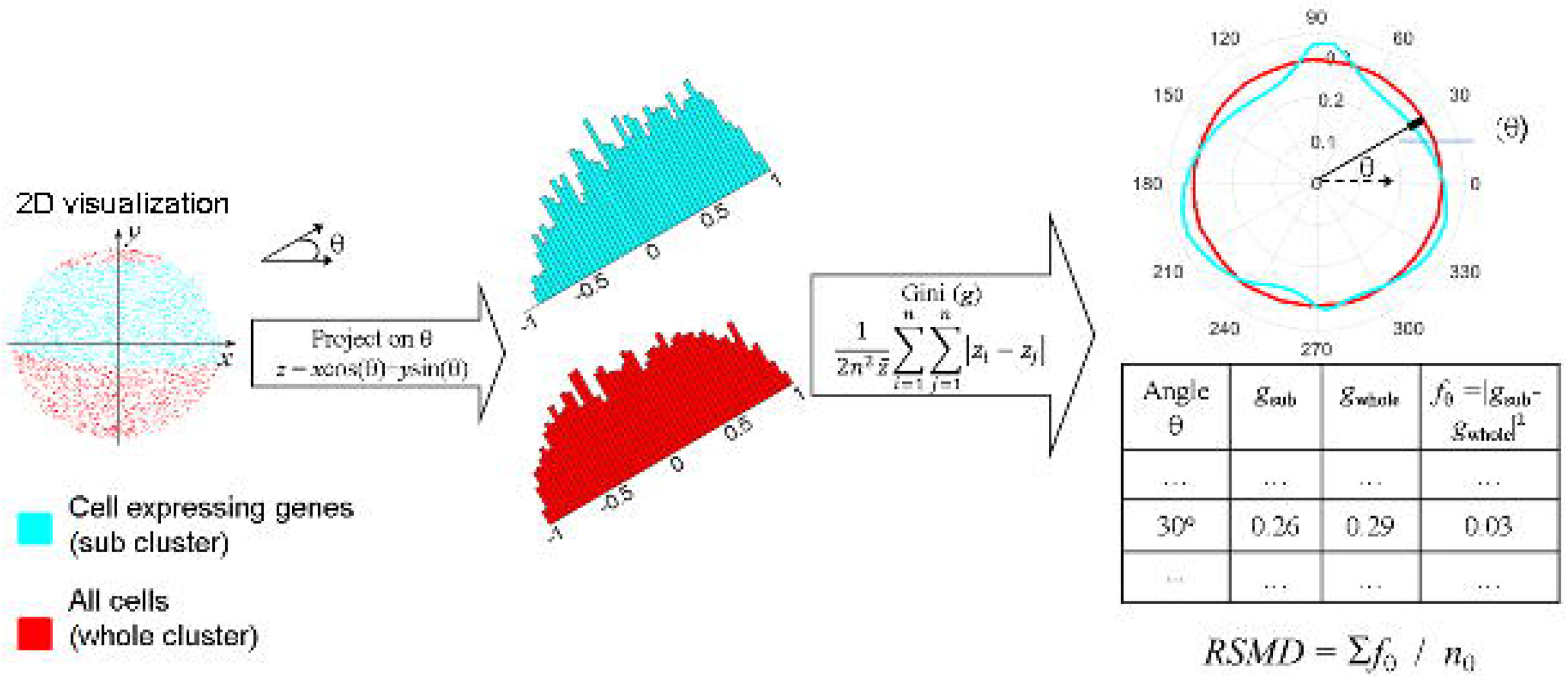
Overall workflow to compute the PGC-RSMD metric for one gene in one cluster of cells. Here, the data points, histogram, and PGC for cells expressing the gene are cyan. The ones for the whole cells in the cluster are red.

We subtracted the scores from (1) with the smallest *z* to ensure that all *z* scores are non-negative, which is the requirement for computing the Gini coefficient. Then, it computed two Gini coefficients *g*_sub_ and *g*_whole_ to measure the inequality among the *z* scores. The *g*_sub_ coefficient only used the distribution of *z* scores obtained from cells expressing the gene. The *g*_whole_ coefficient would use the distribution of all *z* scores. The Gini coefficient formula is as in [28]

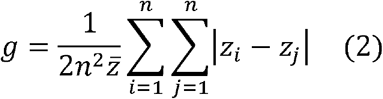

Here, *i* and *j* are arbitrary indices in the list of *z* scores being used in the computation, *z̄* is the average of these *z* scores, and *n* is the size of the *z* score list. Repeating (1) and (2) for multiple angles θ spanning from 0 to 2π would yield the corresponding lists between *g* and θ, as shown in the bottom-right table in **Figure 1**. This would lead to two polar curves for Gini coefficients, one for the cell-points expressing the gene in the cluster, and one for all cell-points in the cluster. We hypothesize that *the two curves would be closer in the marker-gene scenario than in the non-marker gene scenario*. Therefore, we used the root-mean-square deviation (RSMD) metric, which is popular in computing fitness in Bioinformatics [31], to determine whether a gene is a marker in the cluster.

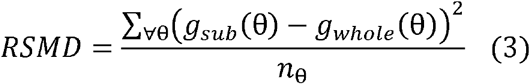

Here, *n*_θ_, also called resolution, is the number of angles θ for which we repeat (1) and (2). In this work, we chose *n*_θ_ = 1000, which makes the angle list θ = 0, π/500, 2π/500,…,999π/500, 2π.

To compute the RSMD statistical p-value for each gene in each cluster, first, we linearly normalized (scaled) the RSMD computed in (3) such that the normalized RSMD is between 0 and 1. This could be done by diving (3) by the largest RSMD among all genes in each cluster. Then, we applied the estimated p-value calculation in [32] to assign a p-value for each gene in each cluster. Briefly, from the RSMD scores in (3), we verified that the RSMD scores followed a bell-shaped distribution. Then, we computed the mean *μ* and standard deviation σ of the normalized RSMD. Then, the p-value for each gene in the cluster is

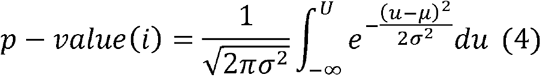

In (4), *U* stands for the normalized RSMD.

### Setting up simulation

In this work, to demonstrate how the PGC-RSMD functions, we setup two simulations. In the first simulation, the cell cluster in the *x*−*y* embedding space had 5000 points, which were uniformly generated in the unit circle *x*^2^ + *y*^2^ ≤ 1. In the second simulation, the 2D visualization of cell clusters had the shape identical to the real-world cluster obtained from visualizing the mouse fetal lung single-cell data [29] using tSNE [33]. We applied the sampling-by-rejection technique [34] to generate these cluster points as follows. In the first simulation, we randomly generated a point whose coordinates are between −1 and 1 using uniform sampling, then accepted the point if it had *x*^2^ + *y*^2^ ≤ 1. In the second simulation, the random point coordinates were within the cluster coordinate range. We extracted the cluster boundary points, compute the polygon from these boundary points, which allowed deciding whether a point was inside the polygon using Matlab [35, 36]. In each simulation, we randomly chose *m* percentage of points (*m* = 5, 10, 15,…,95) and assumed that they represent the cells expressing gene. For each *m* percentage, we repeat the simulation 1000 times.

In addition, to evaluate how the performance of PGC-RSMD would change in drop-out scenario, we modified the single-cell data simulator in [37] as follow. First, we use [37] default parameters to synthesize 2 clusters such that each cluster has 6000 cells, 250 markers (total 500 cluster markers) and other 4500 genes. For each cluster marker, the average expression fol-change when comparing two clusters was between 4 and 1000. We assigned the drop-out probability for each gene from 0, 5, 10,…,to 45% such that there were 25 markers for each drop-out probability. Then, in each cell, we randomly change the marker expression to 0 according to the markers’ drop-out probability. For each of the 4500 non-cluster markers, the expression in each cell was randomly between 0 and 500. We assigned the sparisity – defined as the percentage of non-expressing cells (0 expression) – for each non-cluster marker from 0, 5, 10,…. to 95%. In each cell, we randomly changed the non-cluster marker to 0 according to its sparsity. We used the AUC metric to evaluate whether the PGC-RSMD score could differentiate the 500 cluster-markers: whether each marker is specific for the first or the second clusters.

### Identifying cardiac muscle cell clusters and marker genes from the neonatal mouse heart single-cell data

We obtained the neonatal mouse heart single-cell case-study from the Mouse Cell Atlas [29]. We processed the data as specified in [29]. After preprocessing, the dataset covered 19,494 genes expression in 5075 cells. We use tSNE [33] (without dimensional reduction) to embed the dataset into the 2D space. We used the density-based clustering algorithm [38] implemented in Matlab [39] to identify 9 cell clusters. In the implementation [39], we chose the clustering parameters epsilon = 4, minpts = 40. There were 788, 397, 2966, 156, 288, 123, 76, 125, 87 cell-points in cluster 1, 2,…,9, correspondingly. There were 69 cell-points for which the algorithm is unable to assign to any clusters (Supplemental Data 3).

We computed the percentage of expressing cells (the naïve approach) and PGC-RSMD for all genes in all clusters. We removed genes expressing in less than 10% of the cluster cells. For comparison, in the naïve approach, in each cluster, we selected the top genes sorted by the highest percentage of expressing cells as the cluster markers. In the PGC-RSMD approach, we selected the smallest-RSMD genes with its p-value < 0.05 as the cluster marker. In this work, we focused on identifying the heart muscle cell clusters and their markers. We manually examine the distribution of cells expressing the well-known heart muscle cell markers: *Myh7*, *Actc1*, and *Tnnt2* [40–47].

### Seting up the re-identifying cluster ID problem

To compare the robustness of our PGC-RSMD markers with other approaches, we setup the re-identifying cluster ID as follow. From the visual coordinates and 9 clusters of 5075 cells in [29], we randomly divided the dataset into the training set (4060 cells – 80%) and the test set (1015 cells – 20%) such that set has samples of all 9 clusters. Using the training set and markers’ expression found by PGC-RSMD, in comparison with other approaches, we applied the neural network algorithm [48] to train models that identify cluster ID. We evaluated these models in the test set and recorded the classification accuracy and area-under-receiving characteristic curve (AUC). Here, we hypothesized that the ‘better’ markers would yield higher classification accuracy and AUC. The other approaches being compared with PGC-RSMD are:

- The baseline approach: in this approach, we would train the classification models using all genes expression.
- The differential expression approach: in this approach, we use Fisher’s exact test [49], which computes the likelihood of a gene being expressed (raw expression > 0) in a cluster, compare to the likelihood of the gene being expressed outside the cluster. In this work, we select the DEG markers in each cluster according to the following criteria: odd ratio > 5 and the percentage of expressing cell (*m*) > 50%.
- The SpatialDE [26] approach: SpatialDE finds the gene with high variance regarding the distribution of ‘point’ on the spatial 2D space. The ‘null’ hypothesis in this approach is the gene distribution in the ‘spatial space’ follows a multivariate normal distribution. The marker is selected if the gene expression distribution is significantly different from the null distribution, recorded in the p-value. In this work, we select the SpatialDE marker according to the following criteria: q-value (adjusted p-value) < 0.05 and percentage of expressing cell (*m*) > 50%. In both the DEG and the SpatialDE approach, we sort the markers according to the decreasing order of *m*. To make a fair comparison, we use the same number of markers, ranging from 5 to 100, found by PGC-RSMD, DEG and SpatialDE to train the classification models.

## Results

### PGC-RSMD strongly correlates to the percentage of expressing cell in a cluster

In **Figure 2**, we show that the fitness between the cluster PGC and the sub-cluster PGC strongly correlates to the percentage of expressing cell-points as the ‘sub-cluster’ *m* in the circle-shaped simulation. In addition, as *m* increases, the RSMD variance decreases. We represented the fitness by the **r**oot-**m**ean-**s**quare **d**eviation (RSMD) as showed in the method section. In this figure, for each *m* (from 5 to 95), we randomly generate 1000 sub-clusters and their PGCs. The detailed result of this simulation could be found at the Supplemental Data 1.

**Figure 2.**
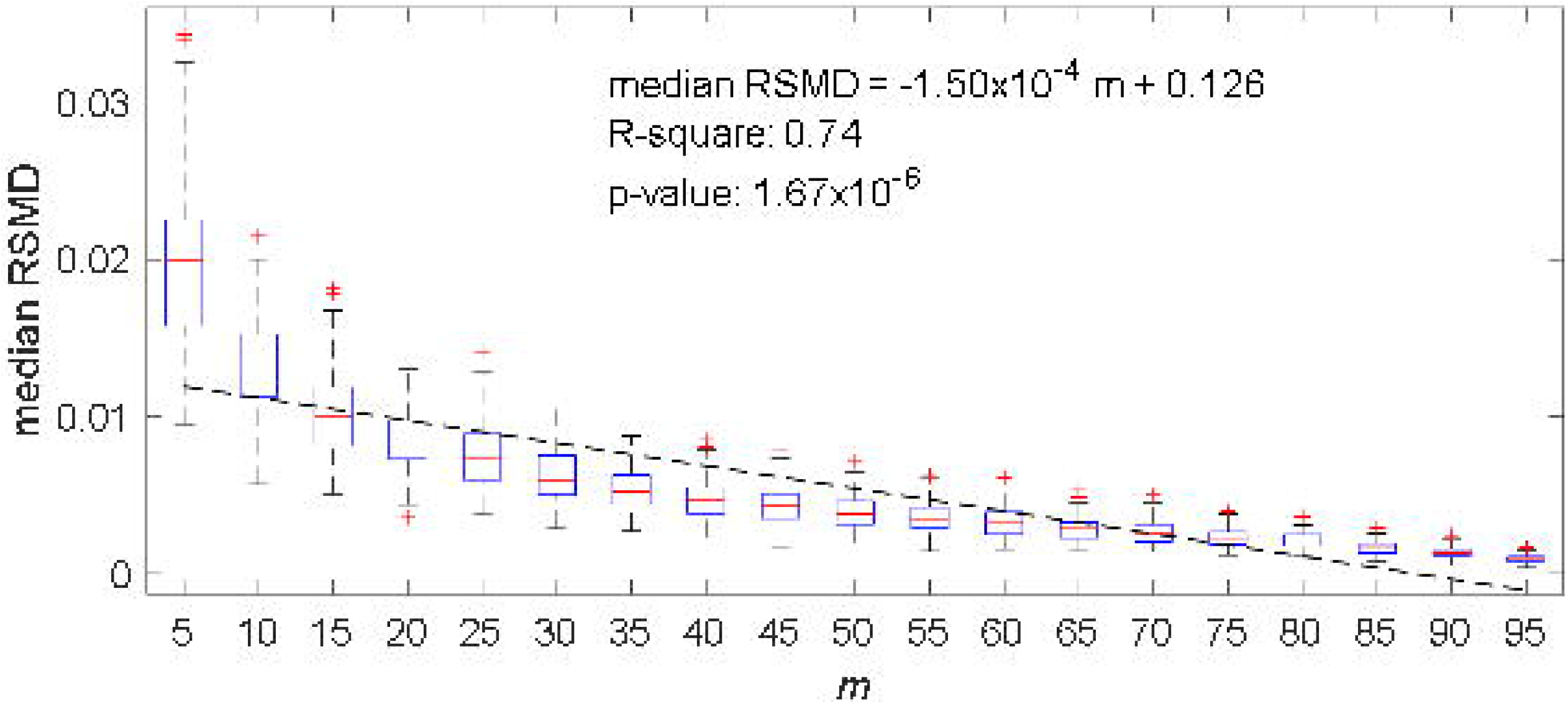
Boxplot showing a strong correlation between ‘subcluster’ percentage (*m*) and cluster-subcluster PGC fitness (RSMD) in uniformly-distributed and a circular cluster.

In addition, we observed a similar correlation when experimenting with the mouse fetal lung single-cell data [29]. **Figure 3a** shows the dataset clusters visualization using tSNE [33] and the chosen cluster. To synthesize a 3000-point cluster with the same shape to the chosen cluster, we still applied the random-by-rejection [34] as presented in the Material and Method section. **Figure 3b** still shows a strong correlation between *m* and RSMD. The detailed result of this simulation could be found at the Supplemental Data 2.

**Figure 3.**
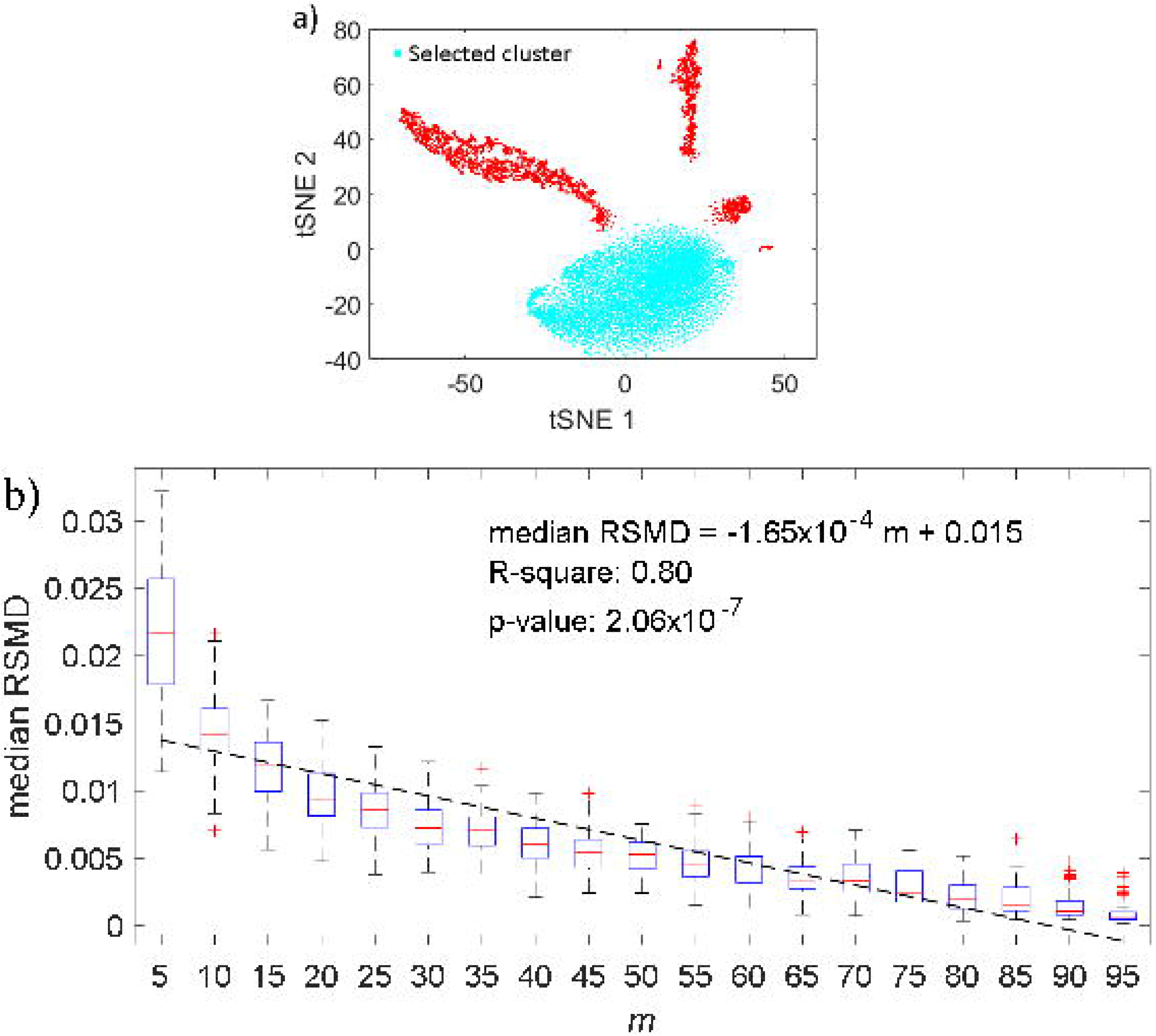
a) The selected cluster for the experiment in [29]. b) Correlation between ‘subcluster’ percentage (*m*) and cluster-subcluster PGC fitness (RSMD) in the selected cluster.

On the other hand, the PGC approach has the potential to answer whether the marker could identify subpopulations of cells in a cluster. **Figure 4a** demonstrates the 30000-point cluster with ring-shape 0.25 ≤ x^2^ + y^2^ ≤ 1, which appears to be a sub-cluster marker. In this case, *m* = 0.75. In this example, RSMD = 0.033 (**Figure 4b**), which is greater than the RSMD distribution computed from the random and uniformly-distributed cluster with the same *m* (**Figure 4c**).

**Figure 4.**
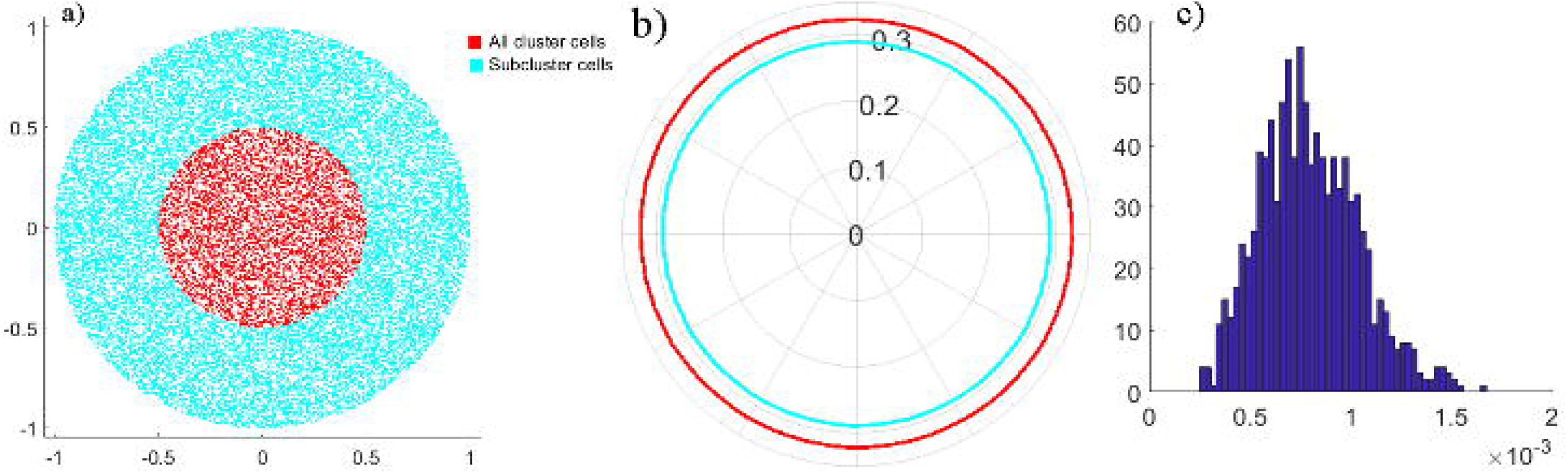
The ring-shape simulation study: a) Visualization of the cluster and ring-shape sub-cluster (*m* = 0.75); b) PGC yield RSMD = 0.033; c) Distribution of RSMD, extracted from Figure 2 with *m* = 75%, when the sub-cluster uniformly distributed on the cluster area.

**Figure 5** shows a decrease of PGC-RSMD performance in the drop-out scenario. Briefly, the synthetic data has 2 clusters, 250 distinct markers for each cluster. Each gene has a specific drop-out rate as presented in the Material and Method section. Using the PGC-RSMD scores in each cluster to differentiate these 500 the cluster-specific markers, we observed that PGC-RSMD achieves very high area-under-receiver-characteristic curve (AUC) (>0.95) when the drop-out probability is small (≤ 5%). However, AUC decreases significantly with the probability of drop-out (**Figure 5a**). This phenomenon further demonstrates the strong association between RSMD and the percentage of expressing-cell. When the drop-out rate increases, the percentage of expressing-cell decreases; therefore, RSMD may mischaracterize a high-dropout marker as non-marker.

**Figure 5.**
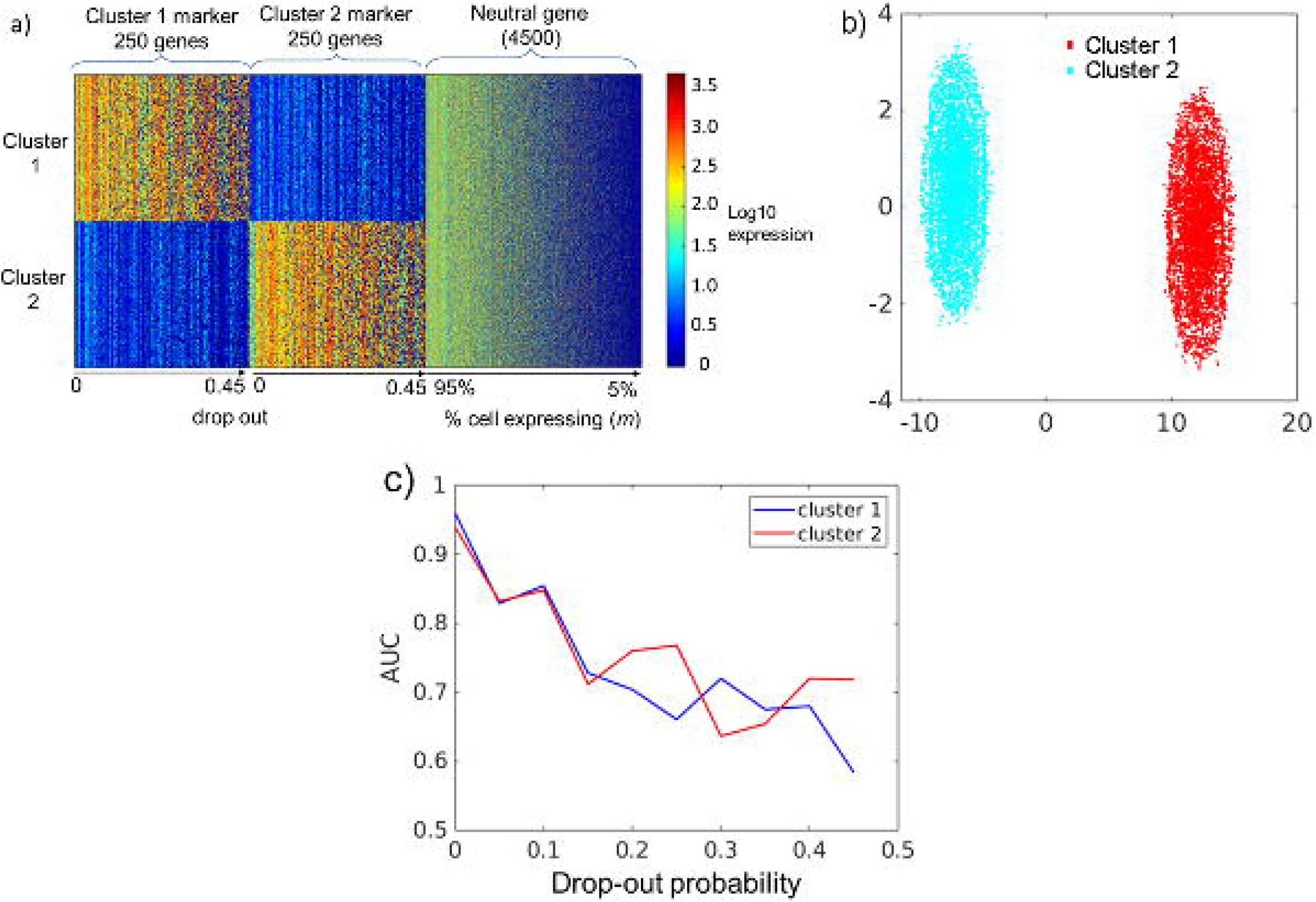
PGC-RSMD performance in recalling cluster marker in drop-out simulation. a) heatmap showing the simulation design of 500 markers and 4500 neutral genes, with drop out / percentage of cell expressing between 5 and 100%; b) The simulation data 2D visualization; c) the AUC drops when drop-out increases.

### Case-study: PGC identifies heart muscle cell in neonatal mouse heart single-cell

#### PGC-RSMD detects markers to support cell-type identification in single-cell mouse neonatal heart data

**Figure 6** summarizes the neonatal mouse heart single-cell data [29] and its 9-cluster markers. **Figure 6a** visualizes these 9 clusters with tSNE. The PGC-RSMD founds 258 genes, which are the union of the smallest 100-PCG-RSMD genes found in each cluster, marking these clusters (Supplemental Data 3). **Figures 6b** and **6c** showed that the gene-cluster marker-association reflects the underlying gene expression in the single-cell data. In these heatmap figures, each row corresponds to one gene.

**Figure 6.**
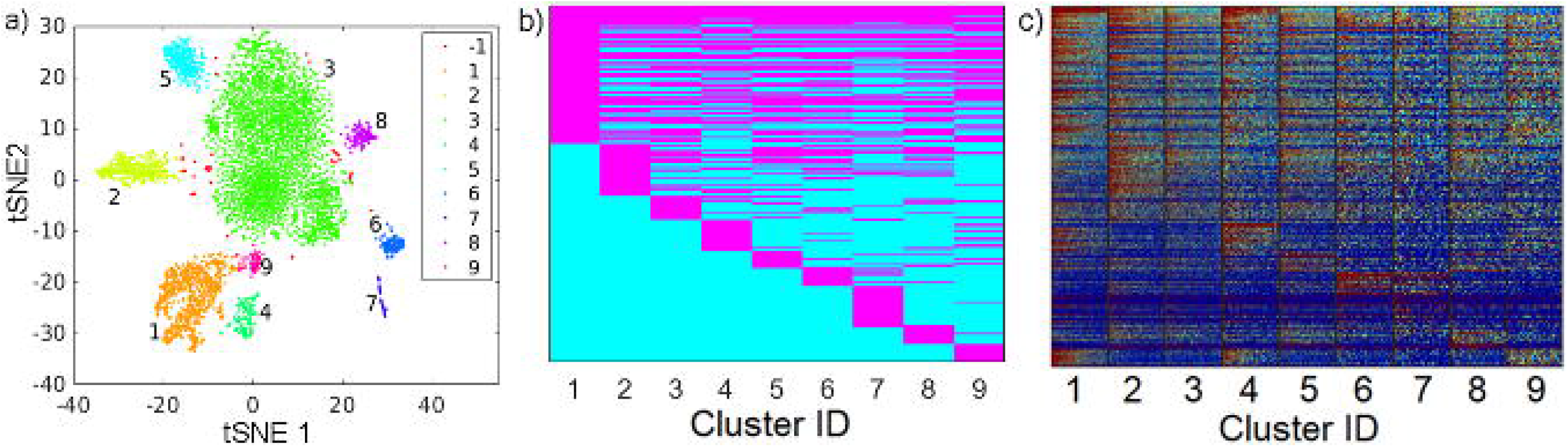
The result from mouse neonatal heart single-cell [29] analysis. a) the tSNE plot shows 9 clusters. b) gene-cluster marker relationship (from 258 genes) found by PGC-RSMD; 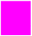 gene is found as marker, 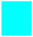 gene is found as non-marker. c) expression heatmap for these genes.

We identified the muscle-cell clusters 1, 4 and 9 by the expression of *Myh7*, *Actc1*, and *Tnnt2*, which strongly express in muscle cell type [40–47] (**Figure 7**). Compared to the naïve method using the percentage of expressing cell, our PGC-RSMD is significantly better by detecting *Actc1*, which are missed by the naïve approach (**Figure 8**). Furthermore, our approach identified *Mgrn1* [50, 51], *Ifitm3* [52], *Myl6b* [53] marking cluster 1, which could play important roles in cardiac muscle functionality, heart failure, and heart development. These genes are not identified using the naïve approach (**Figure 8**). On the other hand, among genes having a high percentage of expressing cell, our PGC-RSMD suggests that *Ndufa4l2*, *Mdh2*, and *Atp5g1* may not be heart muscle cell markers. However, they could suggest a subtype of heart muscle cells (**Figure 9**). The percentage of expressing cells, PGC-RSMD, statistical p-value and ranks for all genes could be found in Supplemental Data 3.

**Figure 7.**
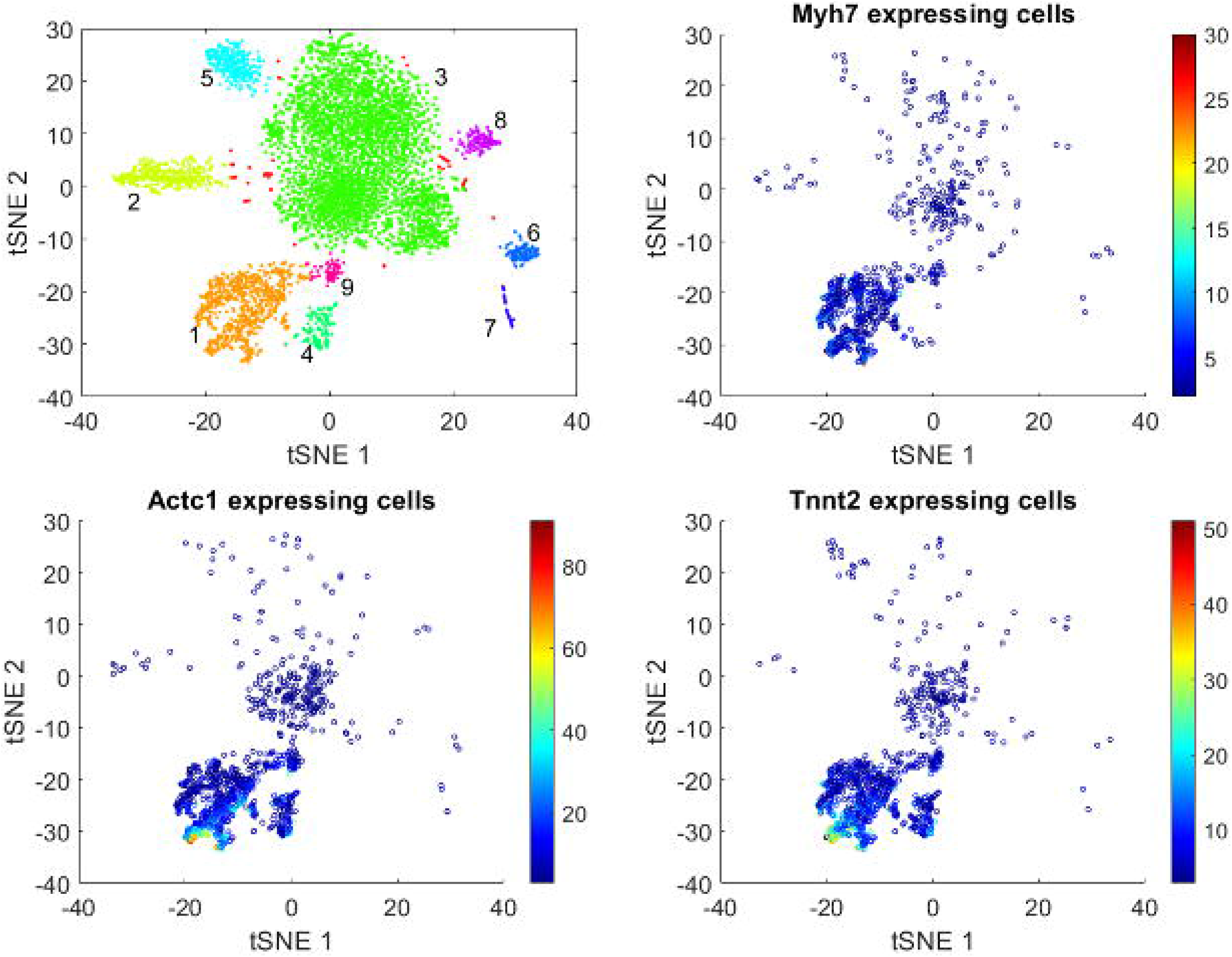
Heart muscle cell clusters, identified by *Myh7*, *Actc1*, and *Tnnt2*

**Figure 8.**
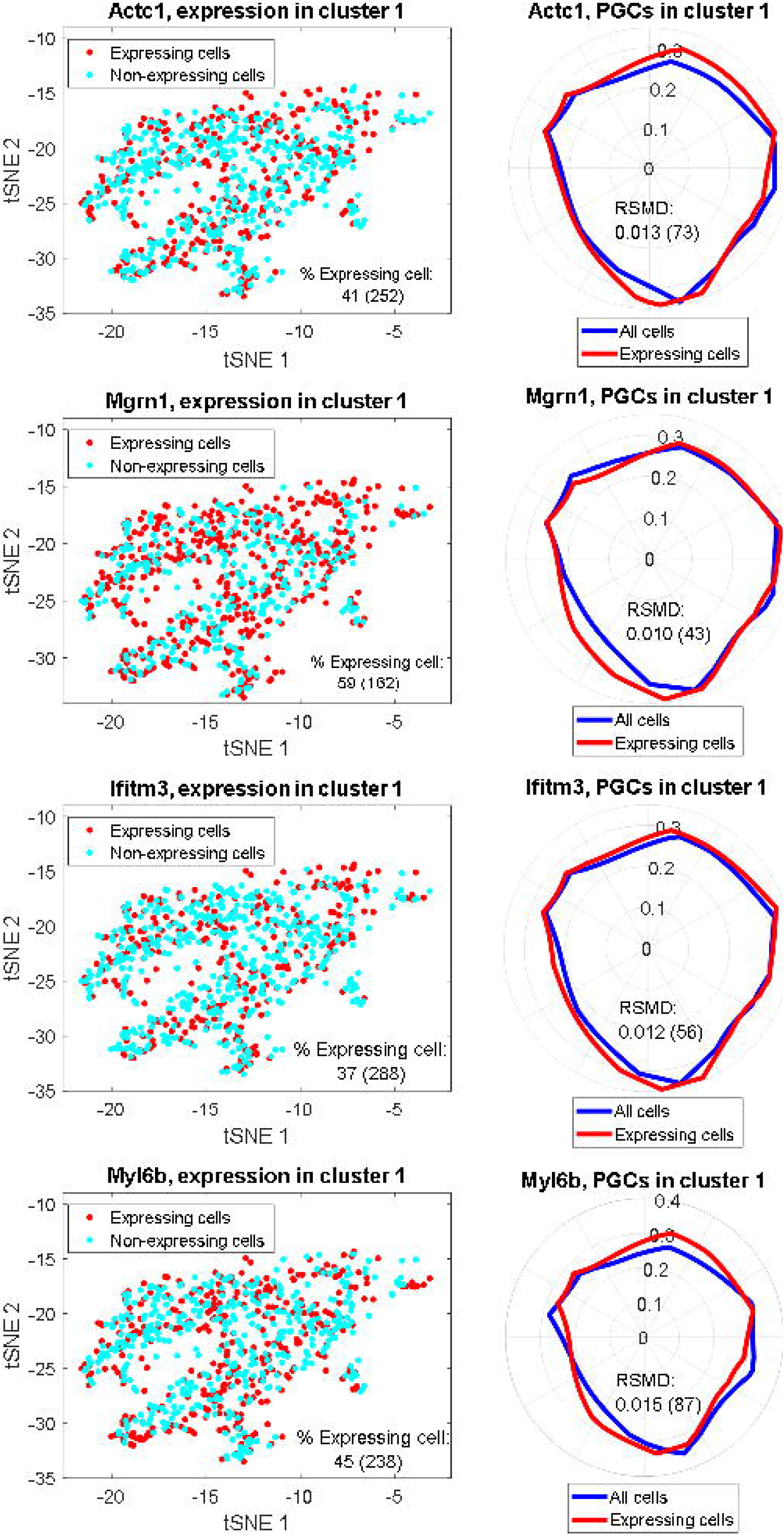
PGC-RSMD highlight makers that do not have high percentage of expressing cells: PGCs of *Actc1*, *Mgrn1*, *Ifitm3*, *Myl6b* in cluster 1. The numbers in the parenthesis are ranks of these genes in each metric

**Figure 9.**
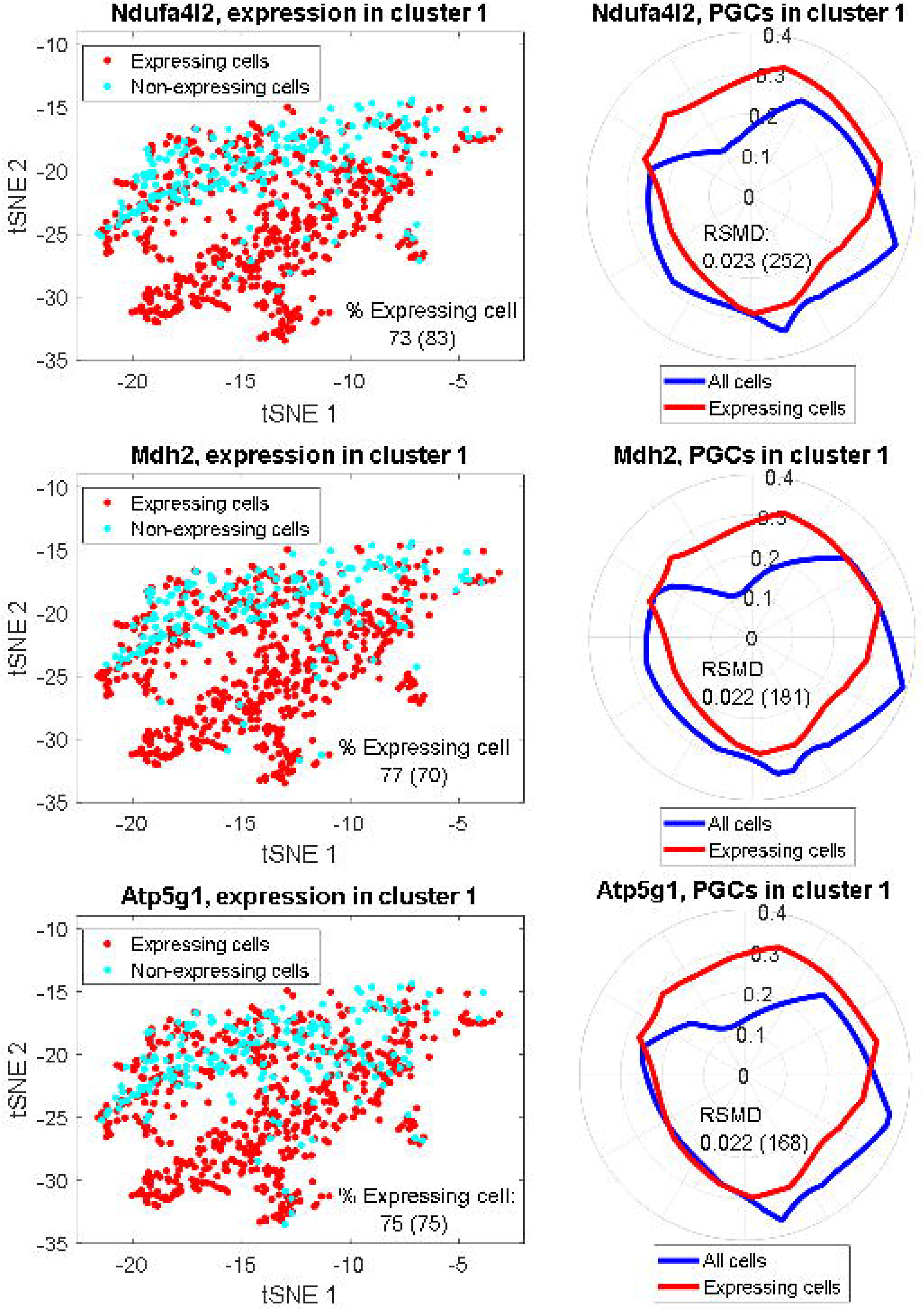
PGC-RSMD shows that gene haves high percentage of expressing cells: *Ndufa4l2*, *Mdh2*, and *Atp5g1*, may not be markers in cluster 1. These genes appear to highlight a local subcluster. The numbers in the parenthesis are ranks of these genes in each metric.

#### Re-identifying the cells’ cluster ID from markers

We observe that the markers found by the PGC-RSMD approach achieve better performance than the similar SpatialDE [26] markers, and similar performance to the differentially-expressed-gene (DEG) when being used to re-identify cell’s cluster ID. Briefly, after computing the visual coordinate and cluster ID of all cells, we randomly split the dataset [29] into the training (80%) and test (20%) sets. We only applied the baseline PGC-RSMD, SpatialDE and DEG approaches to find the markers and built machine learning models to predict the cells’ cluster ID from these markers in the training set. In this experiment, we used all genes to train the predictor in the baseline approach. The detailed description of this experiment could be found in the method section. Evaluating the prediction models in the test set, the PGC-RSMD approach performs closely to the DEG; both have cluster ID prediction accuracy above 0.9 and AUC above 0.95 on average (**Figure 10**). These two approaches significantly outperform SpatialDE, whose accuracy is just above the baseline.

**Figure 10.**
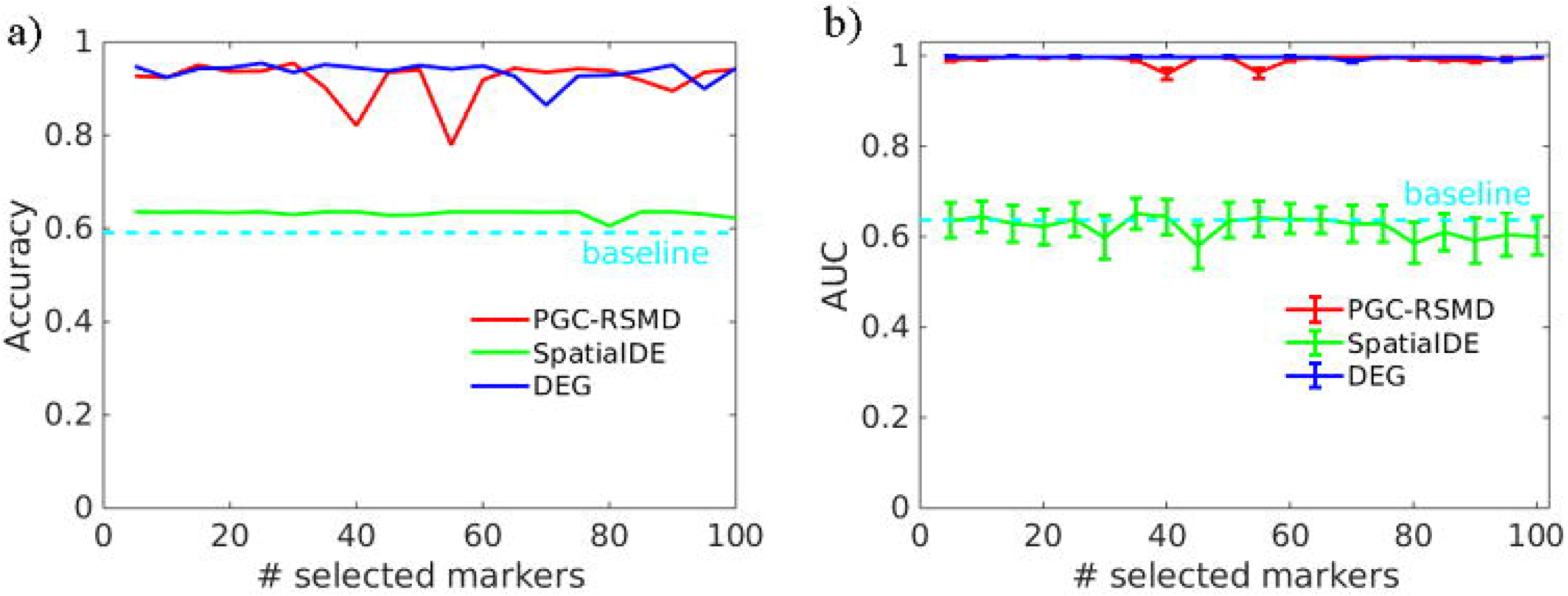
Performance of the PGC-RSMD, SpatialDE, and DEG in re-identifying the cell’s cluster ID problem using dataset [26]. The x-axis shows the number of top-significant markers being selected to train the prediction models. a) accuracy; b) AUC over 9 clusters.

## Discussion

In this work, we show that integrating the embedded information, which does not often have a deterministic relationship with gene expression and is primarily for clustering a visualization, could lead to new insights to biomarkers in single-cell data. In the mouse neonatal heart case-study, our PGC-RSMD approach could recall *Actc1* as the marker characterizing heart muscle cell. Meanwhile, the approach using the ratio of expressing cell may fail to recall because a large percentage of cells does not capture *Actc1* transcript. Therefore, our proposed technique has the potential to handle analytical issues due to single-cell data quality, such as short-read and low sequencing depth [54–56]. On the other hand, for genes having high percentage of expressing cell, the PGC approach could further show that these genes may characterize novel cardiac muscle cell sub-types for future studies, such as in *Mdh2* and *Myl6b*. Therefore, we suggest that the biomarker discovery problem could be divided into two sub-problems: the ‘global markers’ specify cell types and the ‘local markers’ specify subtypes. We could solve these two sub-problems by the right integration of gene expression and visual information.

In this work, we primarily demonstrate how PGC detects markers for single cluster, which does not need the gene expression from other clusters in the dataset. The approach could be extended to incorporate the ‘global’ expression as follow. First, a PGC analysis can be performed with marker cells as the foreground and all cells (regardless of their cluster assignments) as the background. Second, a PGC analysis can be performed for each cluster in the dataset independently and compare among the clusters’ marker lists. In the neonatal mouse heart case study, this approach shows two types of marker: one expressing globally in all clusters, which are likely heart-tissue specific; the other express locally in one or some specific cluster, which are likely cell-type specific.

In addition to our proposed PGC approach, we could apply several alternative strategies to integrate the gene expression and visual information to solve the single-cell biomarker discovery problem. For example, the fractal dimension analysis strategies [57, 58], which focus on evaluating the uniformity of cell-point distribution, could be applied to identify markers in which the expressing cells distribute more densely than they are in the overall cluster. In addition, we could also customize the statistical texture analysis in image processing, such as homogeneity and integrity [59, 60], to analyze the difference between the overall cluster cell-point and cell-expressing gene point as the metric to determine markers. On the other hand, choosing the appropriate visual approach depends on the nature of the data and the problem. Our experiment with the re-identifying cluster ID shows that the well-established SpatialDE [26] does not outperform our approach and the DEG approach. One explanation is that in our problem, a good marker for identifying cell type usually follows a good ‘default’ distribution over the visual space; meanwhile, the SpatialDE aims to find markers that express significantly different from a default distribution.

The major limitation of our proposed PGC-RSMD approach is the long computational time, especially when comparing to the DEG approaches. This is similar to SpatialDE, which also used visual information to detect marker genes. The DEG approaches may only need to compute one statistical test to determine whether a gene is a marker for all clusters. Meanwhile, to draw the curves, PGC-RSMD would need to compute hundreds to thousands, which depends on the desired curve resolution, to characterize one gene in one cluster. Due to the long computational time, we were not able to create multiple simulations, which is the ideal approach, run to compute the statistical [32] p-value for the RSMD score. Therefore, we decided to reapply the estimation presented to compute the p-value. This approach is computationally more efficient but may not well-reflect the statistical characteristic of the single-cell data. In addition, we have not fully tackled the problem of choosing the right threshold to determine whether a gene expresses in a cell. Because of the strong association between PGC-RSMD and the percentage of expressing-cell, we expect that the result would significantly different when choosing a different threshold to determine whether a gene expresses in a cell. In this work, choosing 0 as the threshold still yields good performance because of the high sparsity in the real dataset.

## Conclusions

In this work, we have presented Polar Gini Curve, a novel technique to detect markers from the single-cell RNA expression data using visual information. In principle, our technique could complement the state-of-the-art approach: the PGC technique finds markers such that the expressing cells are evenly distributed throughout the cluster space; meanwhile, the state-of-the-art approach finds markers assuming a multivariate normal distribution of gene expression in the visual space. We have demonstrated that the PGC technique performs better in some tasks in single-cell analysis.

## Authors’ contribution

TN designed and implemented the core polar Gini algorithm, designed the sampling strategies in the simulation, performed the neonatal heart single-cell case-study, and primarily prepared the manuscript. JJ prepared the software package, preprocessed the single-cell data, and executed the simulation designs. NX designed different simulation scenarios, interpreted the statistical outcomes, and prepared the literature review. JC originated the idea of using Gini coefficient curves to integrate gene expression, cell-point distribution, and cluster shape to solve the biomarker discovery problem, designed the performance evaluation, and supervised the overall technical development. All authors reviewed/revised and approved the manuscript.

## Acknowledgments

This work was supported by the ‘startup budget’ granted to Jake Chen from the University of Alabama at Birmingham.

